# Bottlenecks in the transmission of *Porcine reproductive and respiratory syndrome virus* (PRRSV1) to naïve pigs and quasi-species variation during infection in partially immune pigs

**DOI:** 10.1101/320366

**Authors:** Martí Cortey, Gastón Arocena, Emanuela Pileri, Gerard Martín-Valls, Enric Mateu

**Affiliations:** Departament de Sanitat i d’Anatomia Animals, Universitat Autònoma de Barcelona, 08193 Cerdanyola del Vallès, Spain; IRTA, Centre de Recerca en Sanitat Animal (CReSA, IRTA-UAB), Campus de la Universitat Autònoma de Barcelona, 08193 Cerdanyola del Vallès, Spain

**Keywords:** Porcine reproductive and respiratory syndrome virus, PRRSV, deep sequencing, transmission events, founder variants

## Abstract

The existence of bottlenecks during infection of *Porcine reproductive and respiratory syndrome virus* (PRRSV) was studied in an experimental one-to-one model of transmission in pigs. Besides, the differences between viral quasi-species in vaccinated pigs that developed shorter or longer viremias after natural challenge were analysed. The results consistently reported the existence of bottlenecks during transmission. Several positions along the PRRSV genome were identified as being selected in partially immune animals that developed short viremias. Those positions accumulated in GP2, nsp9 and M proteins and resulted in changes in the protein structure and in the interactions of those proteins with their targets. The fact that the affected proteins are known targets of the immunity against PRRSV suggested that the immune response selected those changes. This pig model can be useful for the study of other pathogens of interest in animals and humans.

**Author summary:** Porcine reproductive and respiratory syndrome (PRRS) is one of the most economically important disease of pigs. It is caused by PRRS virus (PRRSV), a positive-sense, single-stranded RNA virus in the *Arteriviridae* family within the order *Nidovirales*. Here, we study the existence of bottlenecks during disease transmission and the differences between viral quasi-species in vaccinated pigs that developed shorter or longer viremias after natural challenge. Our results consistently report the existence of bottlenecks during PRRSV1 transmission and identify several mutations along the viral genome selected by the host immune response that can be clear targets for new vaccine development.

## Introduction

The comprehension of how pathogen transmission occurs is key to understand infectious diseases. Most often, the route of transmission, the portal of entry and the minimum infective dose – when known –, are common features to characterize transmission. However, it is increasingly evident that transmission is an extremely complex phenomenon. For example, because of the existence of bottlenecks during host-to-host transmission [1]. A bottleneck can be defined as a sharp reduction in the size (population bottleneck) or the diversity (genetic bottleneck) in the pathogen population effectively transmitted. The existence of such bottlenecks must be examined considering the portal of entry in the recipient host and the source of the pathogen (blood, nasal secretion, faeces, etc.). Since a pathogen may be present in different tissues, organs, or fluids, each one might be considered a compartment with its own particularities. The pathogen population contained in the compartments where the transmission to the next host occur is termed transmissible population; whereas the successful settlers in the recipient host are called founder variants or transmission founders. The location, size and genetic diversity of the transmissible population can influence the founder population after a transmission event [2–4].

Successful transmission founders can be though as either the result of a non-selective bottleneck – namely, the lucky few that crossed by chance the portal of entry –, or viewed as a selective bottleneck, where only the variants fit enough to cross the portal of entry are transmitted. In both cases, pathogen genetic diversity in the very early phases of the infection would be limited, but with different biological significances. In the case of RNA viruses – that exist as quasi-species – those different scenarios could imply very different outcomes. While the first case – a non-directional unspecific bottleneck – would produce a new quasi-species cloud from randomly selected variants; a directional bottleneck would promote the expansion in the recipient host of variants arising from founders already fit for transmission, although not necessarily the fittest, neither the most efficient for replication in the host.

There are other of factors, such as the immune status of the host that may influence the diversity of a quasi-species. For example, in *Hepatitis C virus* (HCV), continuous diversification has been considered the mean by which the virus escapes the immune system and establishes persistent chronic infection [5]. However, this is an extremely complex phenomenon, where other factors, such as antigenic cooperation between intra-host variants, may permit immune adaptation leading to the co-existence of viral variants with different bind capacities to antibodies, or to be attacked by the cell-mediated immunity [6, 7].

The *ex vivo* study of founder variants and the quasi-species evolution in humans is challenged by the difficulty of determining the precise timing of transmission and the associated quasi-species distribution in the donor. However, the existence of systemic animal diseases caused by RNA viruses with high substitution rates creates an opportunity for examining, in a more controlled environment, transmission bottlenecks and quasi-species variation.

Porcine reproductive and respiratory syndrome (PRRS) is one of the most economically important disease of pigs. It is caused by PRRS virus (PRRSV), a positive-sense, single-stranded RNA virus in the *Arteriviridae* family within the order *Nidovirales*, exhibiting one of the highest substitution rates observed [8, 9]. Experimental models to study PRRSV are well known and have been applied to study transmission [10, 11]. Interestingly, the immune response against PRRSV is unusual, since neutralizing antibodies appear late and cell-mediated immunity has an erratic course for weeks. However, after several weeks of viremia, the virus is confined to the lymphoid tissue and eventually cleared [12, 13]. Pre-formed neutralizing antibodies may protect against the homologous infection in a dose dependent way [14], although that heterologous protection cannot be foreseen [15]. Besides, there is a large individual variation in the immune response [16]. As a result, when a vaccinated animal is challenged, viremia usually develops, but generally of lesser duration than in a naïve animal.

In the present study we used Next Generation Sequencing (NGS) to analyse the quasi-species diversity and evolution in a transmission model of PRRSV in order to: i) characterise and compare the transmissible population and the founder variants in intra-nasally inoculated and naturally infected by contact animals, ii) compare the diversity at early and late phases of viremia and, iii) identify the differences in the viral quasi-species between pigs with partial immunity developing short and long viremias after being in contact with infected pigs.

## Results

### The RNA NGS method was suitable for assessing viral quasi-species

Deep sequencing results for PRRSV1 from sera with high viral load and cell culture supernatants of singly passaged serum samples produced similar viral quasi-species. The estimated error rate between sera and isolated virus ranged between 1 and 3 nucleotides for every 10,000 nucleotides inferred. This rate was considered an acceptable bias. The quality scores (QC) of the NGS runs were above 30 in all the analysed samples, yielding a depth of reads for viral sequences above 115 in all cases. With this depth, variations in the range of percentage units could be determined.

### The characterization of PRRSV transmission events supports the existence of bottlenecks

The transmission experiment scheme and results are summarized in Figure 1. In 8/9 of the intra-nasally inoculated naïve pigs, the viral population in blood at the onset of viremia showed lower diversity compared to the initial inoculum. Similarly, in 4/5 of the naïve pigs infected by directed contact with a seeder, the observed nucleotide diversity was lower compared with their seeder counterparts. As a whole, and assuming the limitations of the method, the results pointed to a reduction in the diversity of the founders compared to the transmissible population, supporting the existence of a bottleneck during PRRSV transmission events.

**Figure 1.**
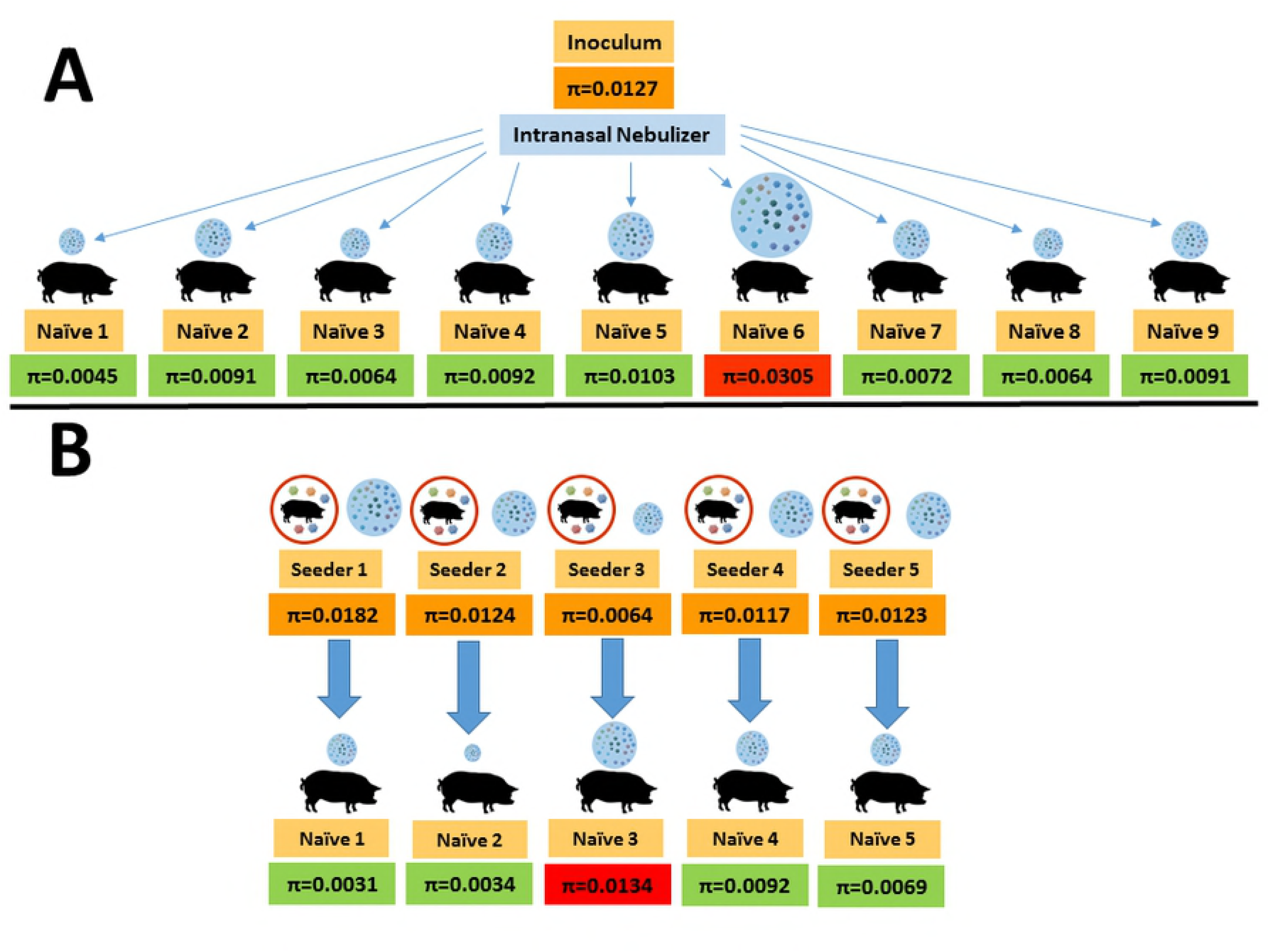
Summary of the 14 transmission events studied: **A**, intranasal inoculation of 9 naïve pigs using a nebulizer; **B**, experimental infection in a 1:1 basis of 5 non-vaccinated naïve pigs from seeders. Nucleotide diversity (π) estimations of the donor population (orange boxes) and the founder variants, within green boxes if the global diversity decreases, in red if an increase is reported.

Taking advantage of the availability of serial samples during the virological course of each animal, it was possible to compare the diversity at different stages (Table 1). After the initial reduction during transmission, nucleotide diversity increased in most pigs analysed, although in some animals this was not the observed pattern.

**Table 1.** Variation of the nucleotide diversity (π) between days of viremia. The table shows the difference in nucleotide diversity between samples of the same animals in different days of the virological course. The difference of nucleotide diversities was calculated by subtracting the π value of a given day from the π value the previous examined day. For the first day of each animal (in red) the difference was calculated with regards to the inoculum (inoculated animals) or with regards to the diversity of the donor on the likely day of transmission (transmission by contact)

| Group | Animal nº | Viremic day | $\pi$ | Difference between consecutive samples |
| --- | --- | --- | --- | --- |
| Inoculum produced in PAM | N.A. |  | 0.0127 |  |
| Inoculated animals | 1 | 1 | 0.0045 | -0.0082 |
|  |  | 6 | 0.0182 | 0.0137 |
|  |  | 13 | 0.0239 | 0.0057 |
|  | 2 | 1 | 0.0091 | -0.0036 |
|  |  | 6 | 0.0124 | 0.0033 |
|  |  | 13 | 0.0115 | -0.0009 |
|  | 3 | 1 | 0.0064 | -0.0063 |
|  |  | 13 | 0.0075 | 0.0011 |
|  | 4 | 6 | 0.0117 | -0.001 |
|  |  | 13 | 0.0528 | 0.0411 |
|  | 5 | 1 | 0.0123 | -0.0004 |
|  |  | 13 | 0.0132 | 0.0009 |
|  | 6 | 1 | 0.0305 | 0.0178 |
|  |  | 6 | 0.0308 | 0.0003 |
|  |  | 15 | 0.0097 | -0.0211 |
|  | 7 | 1 | 0.0072 | -0.0055 |
|  |  | 13 | 0.0524 | 0.0452 |
|  | 8 | 1 | 0.0064 | -0.0063 |
|  |  | 06 | 0.0152 | 0.0088 |
|  |  | 13 | 0.0088 | -0.0064 |
|  | 9 | 1 | 0.0091 | -0.0036 |
|  |  | 6 | 0.0041 | -0.005 |
|  |  | 13 | 0.0335 | 0.0294 |
| Naïve infected by contact | 1 | 1 | 0.0031 | -0.0151 |
|  |  | 6 | 0.0119 | 0.0088 |
|  |  | 22 | 0.0178 | 0.0059 |
|  | 2 | 1 | 0.0108 | -0.009 |
|  |  | 8 | 0.0091 | -0.0017 |
|  |  | 15 | 0.0109 | 0.0018 |
|  | 3 | 3 | 0.0134 | 0.007 |
|  |  | 10 | 0.0165 | -0.0031 |
|  | 4 | 1 | 0.0092 | -0.0025 |
|  |  | 5 | 0.0065 | 0.0027 |
|  | 5 | 3 | 0.0069 | -0.0054 |
|  |  | 10 | 0.0099 | 0.003 |

### Preferred and un-preferred transmitted variants are differentially distributed along the PRRSV genome

Mean frequency differences for each nucleotide in every genome position were calculated for the transmission events where naïve acceptors were infected by seeder pigs (Figure 1B). For this section we used only data from pigs infected by contact, since for the inoculated pigs (Figure 1A), the Porcine Alveolar Macrophage (PAM) propagated inoculum might have shown some level of adaptation to the cell passages and this might have masked the true changes associated with transmission fitness. Figure 2 shows the mean increase or decrease in the frequencies of each nucleotide per position (only values larger than 5% are shown), between the quasi-species of the transmissible population and the transmitted founders. An increase in the nucleotide frequency was associated with a preferentially transmitted variant, while a decrease was identified as an un-preferred transmitted one. Along the PRRSV1 genome, 65 variants in 32 positions were detected (32 preferred and 33 un-preferred). Nineteen nucleotide variants produced synonymous changes, while 13 variants induced potential amino acid changes.

**Figure 2.**
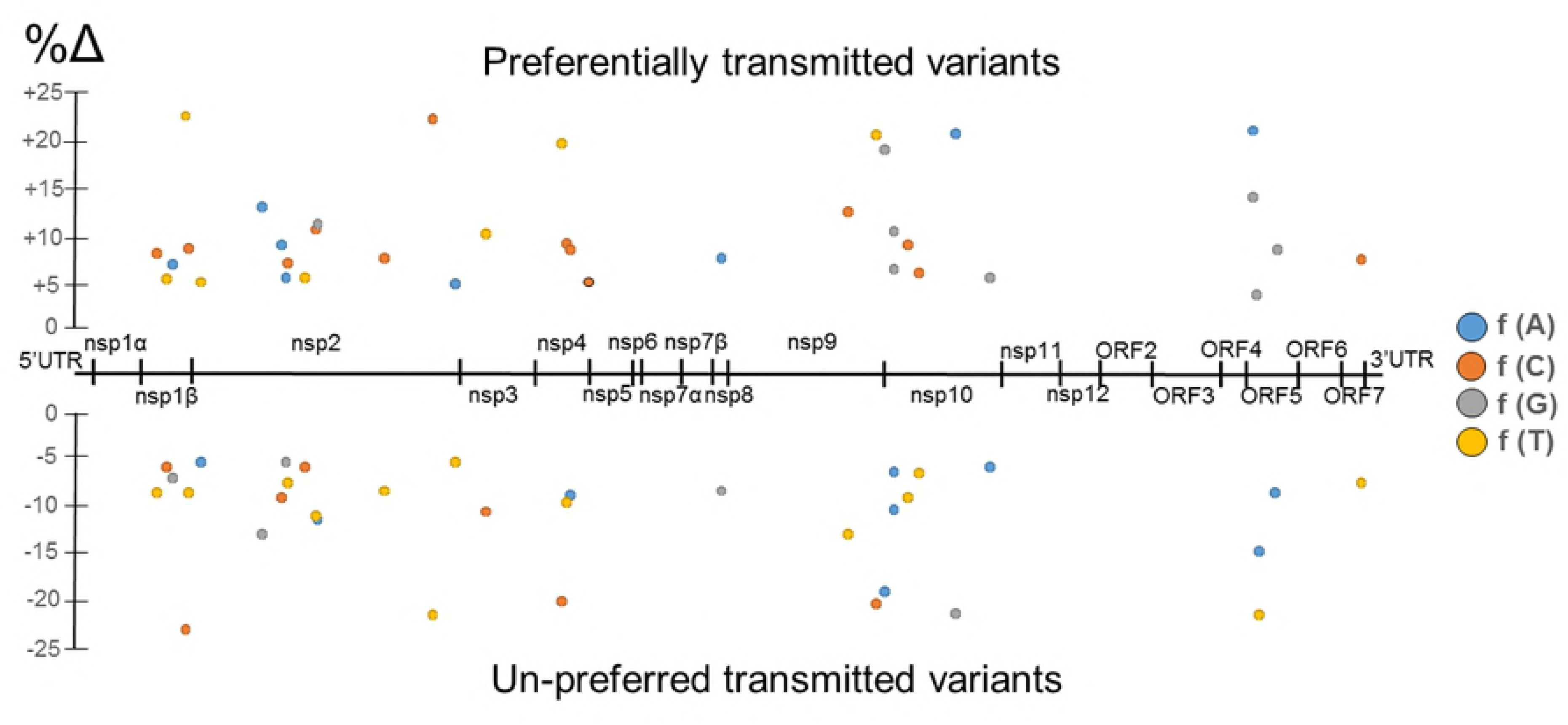
Mean percentage of variation in nucleotide frequencies (only positions with variations larger than 5% are shown), along PRRSV1 genome, of the preferred or un-preferred transmitted variants identified, for the transmission events between infected and naïve pigs studied (Fig. 1B).

A salient feature of the picture was the non-random distribution of the affected positions: 10 in nsp2 region (31%, 4 synonymous and 6 non-synonymous), 3 positions in nsp4 (9%, 2 synonymous and 1 non-synonymous), 3 in nsp9 (9%, all synonymous), 6 in nsp10 (19%, 3 synonymous and 3 non-synonymous) and 3 in ORF5 (9%, 1 synonymous and 2 non-synonymous).

Next, we examined and related the heteronymous changes with the regions and known features of the encoded proteins. The six amino acid changes detected within nsp2 fall in the two hypervariable regions flanking the papain-like cysteine protease domain, with one change (Gln-336-Lys) located in the B-Cell epitope site 3 proposed by [17] and a second one (Tyr-736-His) falling in a non-conserved epitope inducing IFN-γ and IL-10 responses, as reported by [18]. In nsp4, an Ile-142-Leu change is located in the middle β-barrel domain II of the main PRRSV proteinase 3C-like protease, but the change did not result in any substantial change in the 3D structure of the protein. The single change detected in nsp5 – Leu-32-Phe – was located in a transmembrane domain. Regarding nsp10, 2 out of 3 amino acid changes identified in this protein fall in the Zinc-binding domain, while the third was located in the C-terminal domain, downstream the helicase. One of the affected residues, located at position 46, preferentially changed from Ser to Gly during the transmission events. Finally, in GP5 two nucleotide mutations in the first and second position of a codon leaded to the change Ile-36-Asp in the hypervariable ectodomain of GP5, just after a signal peptide cleavage site. None of the abovementioned changes produced a relevant structural change in any protein.

### Differences in viral quasi-species between short- vs long-viremia pigs are mostly located in nsp9 and ORF2

In PRRS, immunity against heterologous viral strains is considered partial and, in consequence, the infection of vaccinated animals is possible if a heterologous strain is used. After examining transmission to naïve pigs, we compared the viral quasi-species in two groups of partially immune pigs infected by contact from seeder penmates in a one-to-one basis. Those partially immune animals developed viremias that were classified as long (*LV*) or short (*SV*). The results indicated a different distribution of the changes in nucleotide frequencies between groups along the viral genome (Figure 3). Forty-five of the 55 positions (81.8%) identified were located in the nsp9 (37 positions, 64%) and ORF2 (8 positions, 15%); while these two proteins only account for 17.8% of the nucleotide positions in the viral genome. Remarkably, in all these 55 positions, the nucleotide variants characterising the *LV* group coincided with the nucleotide present in the original strain causing the infection, while the *SV* group always showed a different nucleotide. When the nucleotide variants present in the *SV* and *LV* were translated, most variants (43) generated the same amino acid (blue dots in Figure 3), but 12 introduced amino acid changes: 2 located in nsp2, 5 in nsp9, 3 in ORF2 and 2 in ORF6 (red dots in Figure 3).

**Figure 3.**
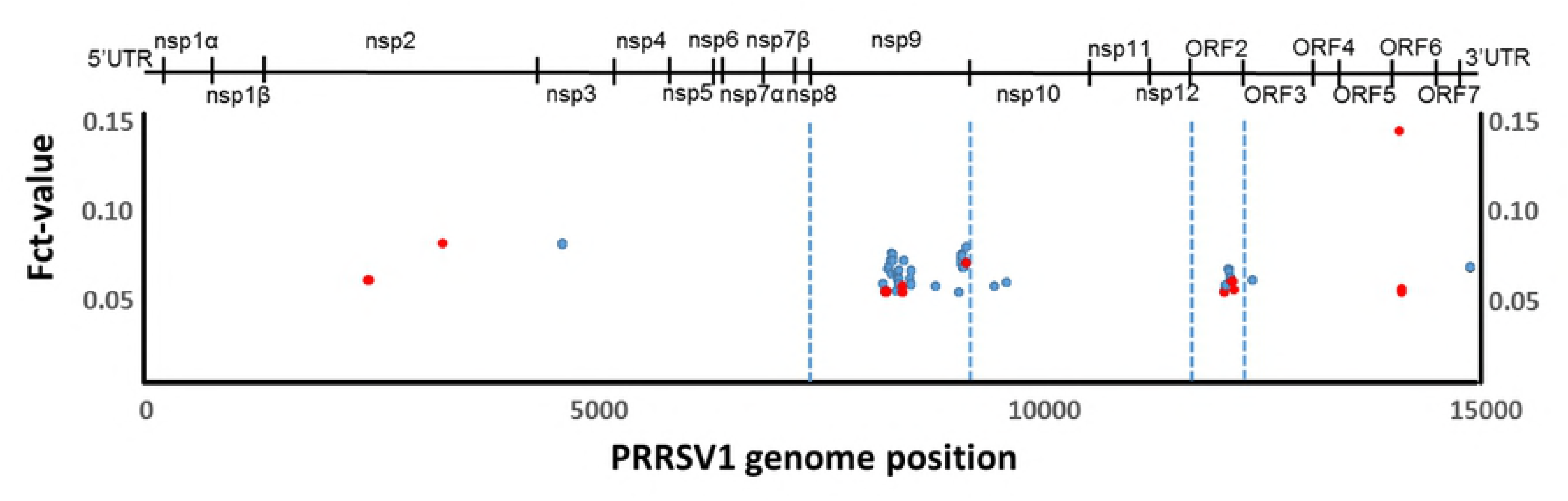
Location of the nucleotide positions along the PRRSV1 genome showing differences (calculated as Fct results, only values >0.05 are shown) between two groups of vaccinated animals developing short- (n=7) or long-viremia (n=4) against PRRSV1 infection.

Again, non-synonymous changes were examined for potential biological significance. The two changes identified in nsp2, Ala-392-Thr and Pro-669-Ser, fall in the B-cell epitopes 4 and 7 described by [17]. In nsp9, one of the key enzymes for PRRSV synthesis, all the amino acid changes identified were located in the viral RNA-dependent RNA polymerase (RdRp), encoded by the C-terminal domain. The comparison of the RdRp 3D structures between *SV* and *LV* groups (inferred with SWISS-MODEL), indicated that the amino acid changes Lys-338-Arg and Lys-641-Arg do not produce any evident structural change. On the contrary, the change Gly-387-His, caused by nucleotide changes in the three positions of the codon, induced a structural change. The presence of a 387Gly in *SV* group resulted in the formation of an α-helix between residues 384–388 and reduced the positive charge at the opposite side of the active centre of the protein where Mg^2+^ molecules bind. (Figure 4).

**Figure 4.**
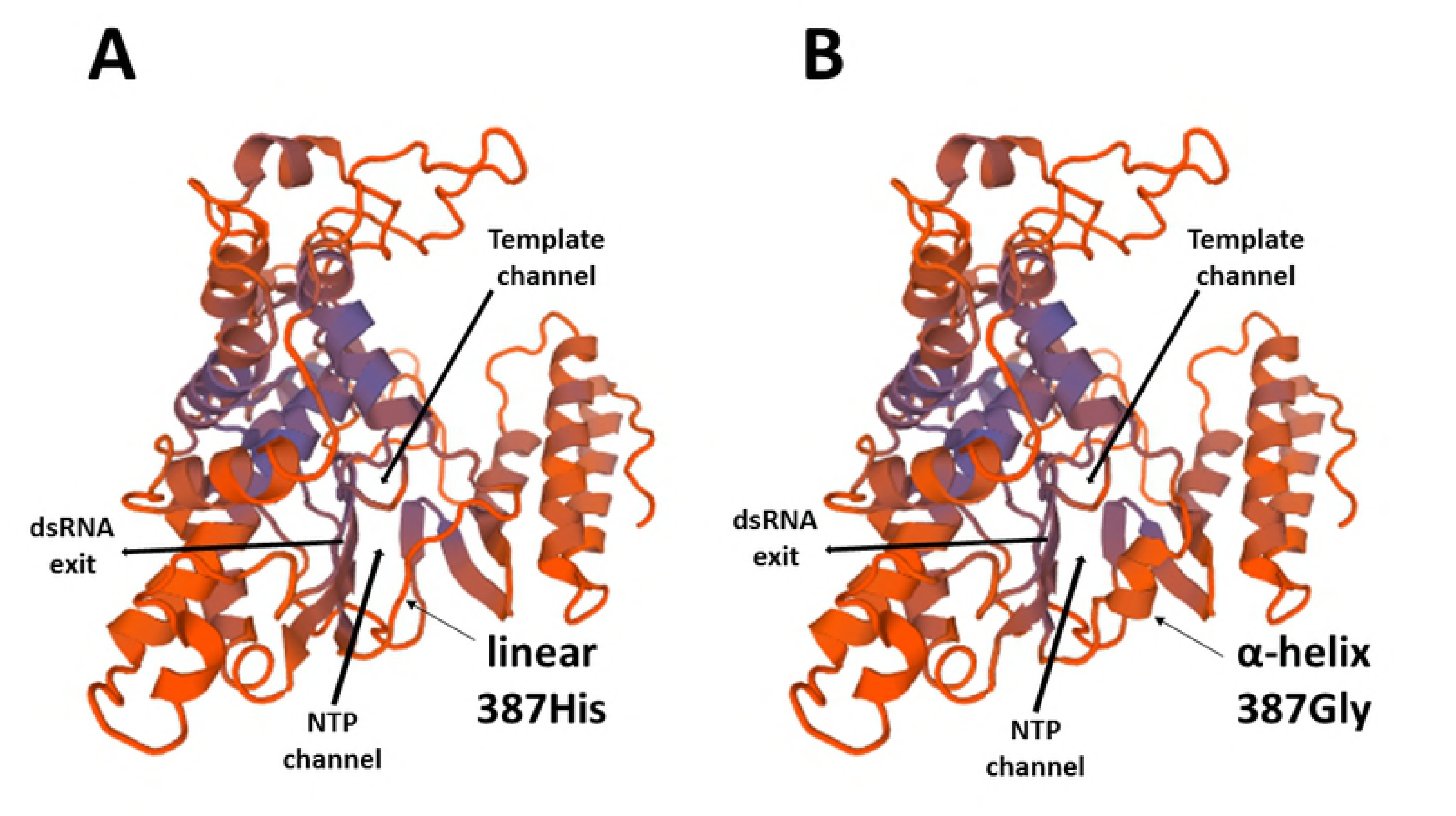
3D-structures of the PRRSV1 RdRp based on the translated sequences of nsp9 from the viral quasi-species of the inoculum (A), long- (A) and short- (B) viremia pigs. The NTP and the template channels, the dsRNA exit, as well as the amino acid residue 387 are indicated.

In GP2, the three amino acid changes identified, namely Gln-146-Arg, Val-151-Ala and His-184-Arg. fall in the ectodomain, slightly downstream the linear B-cell epitopes proposed by [19] and [20]. The amino acid changes reported for this protein implied more positively charged residues in *LV* compared to *SV* group.

Finally, the last two amino acid changes, Asp-10-Asn and Stop-25-Tyr, are located in the short stretch exposed at the virion surface in the N-terminal ectodomain of protein M, encoded by ORF6. The Asp-10-Asn imply a charge modification from neutral (*LV*) to negative (*SV*).

## Discussion

With the introduction of NGS technologies, the experimental analysis of viral genetic diversity has changed dramatically. Due to its massively parallel approach, NGS generates millions of reads that cover every nucleotide position from thousands to hundreds of thousands times. Hence, low-frequency variants within viral quasi-species can be adequately detected and characterized [21] and it is possible to see variations that, although significant, would remain unnoticed using Sanger sequencing. However, the usefulness of NGS for viral diversity estimations depends crucially on the quality of the sample and on the procedure to prepare it [22]. In the present work we applied a tailor-made NGS method to characterize the diversity of PRRSV quasi-species. The method omitted the PCR amplification step and used instead a singly passage in PAMs when needed because of the low amount of virus present in many samples. By using PAMs, the same cells that support viral replication in the host, a low bias was generated in a single passage and the error rate produced was also very low. This approach could be useful for other viruses that cannot be analysed directly by NGS from biological samples because of the low titres present.

As stated, virus populations may face bottlenecks during the infection cycle [1]. When transmission takes place through a mucosal portal of entry, infection is usually initiated by a limited number of viral particles compared to the total number of particles reaching that portal [4]. Besides population bottlenecks, there are many examples of extreme genetic bottlenecks in RNA viruses such as *Human Immunodeficiency Virus* (see reviews by [4] and [23]), HCV [24–26] and *Simian immunodeficiency virus* [27].

In most of the experimentally inoculated pigs and the animals infected in a quasi-natural way by contact with infected seeder pigs, viral diversity was reduced during transmission, supporting the existence of a bottleneck during PRRSV infection. The nature of such a bottleneck is more difficult to precise. The changes in the nucleotide diversity were not scattered randomly across the viral genome, but focused in a few targets. Beforehand, one could think that the structural proteins interacting with the target cells would be the most affected ones, since they are the first interacting with the mucosa surface. Interestingly, this was the case with GP5, the viral glycoprotein establishing the first interaction with porcine sialoadhesin, one of the viral receptors on the macrophage surface [28]. The preferentially transmitted variant introduced a change in position 36 favouring an Asp, a more acidic amino acid. It is difficult to interpret this result but it is located close to a potential glycosylation site and adjacent to the neutralization epitope in GP5 [29]. It is tempting to hypothesize a potential increased interaction between GP5 and siloadhesin favoured by the higher polarity of the 36Asp variant. Besides, it is worth noting that all but one of the other favoured changes affected non-structural proteins, pointing towards the selection of variants with different replication characteristics, although the result of the precise amino acid changes could not be ascertained from the literature.

After the initial diversity reduction during the transmission event, the viral diversity of the circulating quasi-species increased in most cases, as expected in an initial expansion of a viral population in a naïve animal. Afterwards, in later stages, diversity could increase or fall but since a detailed characterisation of the immune response at each time point was not performed, the causes are not clear.

The third objective of this work was to analyse the quasi-species differences between two groups of partially immune pigs developing short and long viremia. In the present work, the duration of the viremia was correlated with the titers of neutralizing antibodies against the virus [10]. Therefore, beforehand, main changes were expected to be located in potential targets for the antibodies. Neutralizing antibodies for PRRSV have been reported to be induced by GP2, GP3, GP4, GP5 and M proteins, although immune-dominant epitopes are mainly located in N and nsp2 [14, 30]. In the present case most of the changes in *SV* occurred in nsp9, followed by ORF2 that encodes GP2 (Figure 3). The 3D modelling of nsp9 (RdRp, Figure 4) showed that the introduction of a Gly-387-His after changes in the three nucleotides of the codon, produced a change in the folding of the protein, from linear to α-helix. This change resulted in the absence of a positively charged group opposite to the site of union of Mg^2+^, and would probably cause a modification in the electrostatic forces involved in the interaction with the NTP channel. It is reasonable to think that such a change would affect the efficiency of RNA synthesis.

Regarding the changes in the ORF2, the variants found in *SV* also resulted in a less charged protein. GP2 is known to interact together with GP3 and GP4 with CD163 [31], the essential cell receptor for PRRSV [32, 33]. Again, this could induce a weaker interaction with the receptor CD163.

Apart from the abovementioned changes in nsp9 and ORF2, an additional interesting change was observed in the M protein (encoded by ORF6). The negative charge present in Asp10 (*SV*) may interfere in the disulphide link between the residue 9 of M protein and the residue 48 of GP5 [34]. Therefore, potential weaker interactions in the variants present in *SV* pigs with heparan-sulphate, another receptor protein for PRRSV may be induced [35].

In the frame of animals with higher titers of neutralizing antibodies, as observed in the *SV* group [10], the changes reported in GP2 and M could be understood as escape mutations. The lower antibody titers in the *LV* group probably prevented those changes from being positively selected, and the major variants present in the initial inoculum remained the commonest. Similarly, it was proposed that T-cell responses contributed to partial levels of cross-protection in PRRSV, and therefore, the changes observed in nsp9 could be seen as escape variants [36]. Potential T-cell epitopes for PRRSV have been proposed for nsp9 [37], as well as for the RdRp of other *Nidovirales* [38, 39]. The striking fact relates with the fact that those changes in nsp9, GP2 and M would probably result in less efficient viral variants for, either interacting with the cell receptor, or replicating. This scenario suggests that in animals with higher levels of immunity, the variants escaping the immune system are not fit enough to maintain a viable quasi-species cloud, and consequently, the viral population collapses. Accordingly, nsp9, GP2 and M would be clear targets for new vaccine development.

In summary, the present report shows a feasible approach to study transmission events and changes in viral quasi-species during the course of an infection in pigs. The results were compatible with the existence of a transmission bottleneck for PRRSV and showed some targets for understanding the effects of the immune response on viral diversity. This pig model could be used to study human diseases such as influenza and to gain understanding of how transmission of RNA virus occur.

## Materials and Methods

### Animal experiment

Samples used in the present study were obtained in the course of a previous experiment aimed to determine the transmission of PRSRV in a one-to-one basis [10]. Figure 1 summarizes the first experiment. Two different scenarios were examined in the first experiment regarding transmission. The first one studied animals experimentally infected by the intranasal route with a PRRSV inoculum produced in PAMs (n=9); the second scenario consisted in cases of transmission by contact to naïve pigs (n=5). Those PAMs were obtained from bronco-alveolar washes of lungs collected from four-week old high health pigs. In all cases, samples used were sera collected in the first observed day of viremia for the recipient, usually day 2 after inoculation or contact and when transmission occurred by contact, the closest day of viremia to that event for the donor (usually 1–3 days before the onset of the viremia in the recipient). Additionally, since samples at later viremia stages were available we examined the changes in the diversity throughout the viremic period.

In a second experiment, the changes in the viral quasi-species present in blood of partially immune animals (previously vaccinated), that got the infection by contact with seeder pigs was examined (n=11). For this purpose, the last day of viremia was analysed. In this later experiment, animals were seen to develop long viremias (≥7 days), or short viremias (<7 days) and accordingly, were classified in two groups: short viremia (*SV*, n=7) or long viremia (*LV*, n=4). Since in some cases the amount of virus in blood was not enough to proceed directly to NGS, all samples were subjected to a single passage in PAMs in order to maintain similar conditions for all.

For the first experiment, the inoculum used was a sixth passage of strain CReSA3267 (Accession Number JF276435). In both experiments, viral load in blood was determined by one step RT-PCR (qRT-PCR) targeting PRRSV ORF7 using the method described by [40].

### NGS protocol

Cell culture supernatants were firstly centrifuged up to 14,000 g in order to remove potential debris. Afterwards, total RNA was extracted using the Trizol LS© reagent according to the manufacturer directions. Extracted RNA was assessed by spectrophotometry at 260 nm and 280 nm and used in the PRRSV-specific RT-PCR as above for determining concentration of RNA and purity.

The assessment of PRRSV diversity within each serum sample was characterised directly from RNA without any previous amplification step using a NGS approach developed by our group. The procedures included: i) the construction of a genomic library for Illumina NGS sequencing using a commercial protocol and reagents (Protocol for use with Purified mRNA or rRNA Depleted RNA and NEBNext^®^ Ultra™ II RNA Library Prep Kit for Illumina^®^, New England Biolabs), ii) trimming of low quality reads (QC>20 as determined by FastQC©software, Babraham informatics) using Trimmomatic© [41], iii) mapping of the reads against strain CreSA3267 using the Burrows-Wheeler Aligner applying the BWA-MEM algorithm for long reads [42], iv) variant calling with SnpSift© to determine the frequency of each nucleotide at each position of the reference genome and, v) construction of the viral quasi-species in fasta format.

### Validation of the procedure, estimation of the PAM passage error rate and quality control check

Given that the samples used were cell culture supernatants, it was necessary to validate the technique for ascertaining the potential bias introduced by the single passage in PAM. For this purpose, four samples with an estimated viral load of >10E6 viral genomes/ml (after quantification by RT-PCR) were directly used in the abovementioned protocol. In parallel the same samples were singly passaged in PAM and the cell culture supernatants were examined similarly. To be subjected to further analysis, a sample should produce a complete genome with at least a depth of 100 reads per position.

### Nucleotide diversity estimations and frequency changes per position in the transmission events

For assessing the change in the diversity of viral populations in the different transmission cases, the calculation of the nucleotide diversity (π) was performed using DNAsp [43]. These calculations were comparing respectively: a) the diversity in the PAM inoculum versus the diversity in the samples collected the first day of viremia in experimentally inoculated animals and, b) the diversity in the first day of viremia of animals infected by contact versus the diversity in the sample of the donor the most likely day of transmission. Additionally, for the transmission by contact cases (5 animals) the frequency of each nucleotide in each position in the donor and the recipient were compared. With this it could be estimated what nucleotides increased or decreased their frequency in the transmission event. Frequency changes above 4.9% were arbitrarily considered relevant.

### Analysis of molecular variance (AMOVA) in partially immune animals

In the case of partially immune animals, it was considered key to identify nucleotide positions that could be differentially selected in animals with longer or shorter viremias. Therefore, firstly animals were grouped according to the viremia as stated before. Then, the average frequencies of each nucleotide per position and group were compared using AMOVA in Arlequin ver 3.5.2.2 [44]. Positions presenting a Fct>0.05 were considered for further analysis.

### Screening of potential changes in the amino acid composition in the viral proteins

For all cases (transmission to naïve or to immune animals), the resulting reads were ordered to represent the different viral proteins known. Once this was done, for the positions showing changes above the considered threshold for the frequencies of the nucleotides, the potential changes in codons were analysed.

In a first step, each change potentially causing the arising or the increasing of a heteronymous codon was annotated in the corresponding domain of the protein if known. Then, by bibliographic review, the affected positions were assessed for a known function. In addition, the changes in the protein structure were evaluated using SWISS-MODEL (https://swissmodel.expasy.org/repository/) and the potential modification in the charge of the protein or site were evaluated.

